# Gene-flow from steppe individuals into Cucuteni-Trypillia associated populations indicates long-standing contacts and gradual admixture

**DOI:** 10.1101/849422

**Authors:** Alexander Immel, Stanislav Țerna, Angela Simalcsik, Julian Susat, Oleg Šarov, Ghenadie Sîrbu, Robert Hofmann, Johannes Müller, Almut Nebel, Ben Krause-Kyora

**Affiliations:** Institute of Clinical Molecular Biology, Kiel University, Kiel, Germany; “High Anthropological School” University, Chişinău, Republic of Moldova; “Olga Necrasov” Centre for Anthropological Research, Iași, Romania; Institute for the History of Material Culture, Russian Academy of Sciences, Saint Petersburg, Russian Federation; Institute of Cultural Heritage, Academy of Sciences of the Republic of Moldova, Chișinău, Republic of Moldova; Institute of Prehistoric and Protohistoric Archaeology, Kiel University, Kiel, Germany

## Abstract

The Cucuteni-Trypillia complex (CTC) flourished in eastern Europe for over two millennia (5100 – 2800 BCE) from the end of the Neolithic to the Early Bronze Age. Its vast distribution area encompassed modern-day eastern Romania, Moldova and western/central Ukraine. Due to a lack of existing burials throughout most of this time, only little is known about of the people associated with this complex and their genetic composition. Here, we present genome-wide data generated from the skeletal remains of four females that were excavated from two Late CTC sites in Moldova (3500 – 3100 BCE). All individuals carried a large Neolithic-derived ancestry component and were genetically more closely related to Linear Pottery than to Anatolian farmers. Three of the specimens also showed considerable amounts of steppe-related ancestry, suggesting influx into the CTC gene-pool from people affiliated with, for instance, the Ukraine Mesolithic. The latter scenario is supported by archaeological evidence. Taken together, our results confirm that the steppe component had arrived in eastern Europe farming communities maybe as early as 3500 BCE. In addition, they are in agreement with the hypothesis of ongoing contacts and gradual admixture between incoming steppe and local western populations.

## Introduction

In the archaeological record of eastern Europe, the first evidence for an agrarian lifestyle appeared in the 6th millennium BCE, when the Neolithic societies of the Danube basin (e.g. Linear Pottery [Linearbandkeramik, LBK] and Starčevo) began to spread to the Carpathian region^1, 2, 3^. Following these early foundations, a new society, the Cucuteni-Trypillia complex (CTC) emerged in a vast area that encompassed modern-day eastern Romania, Moldova and western/central Ukraine (Trypillia; Figure 1). CTC flourished in eastern Europe for about over two millennia (5100 – 2800 BCE) from the end of the Neolithic to the to the Early Bronze Age, and is commonly divided into an Early, Middle and Late period^4, 5^. Due to its geographic location, CTC was at the nexus of several contemporaneous societies, such as the Lengyel, Funnel Beaker (FBC, also Trichterbecher TRB) and the Globular Amphora (GAC) cultures (Figure 1)^6^. CTC is characterized by a wealth of material finds, attesting to a strong farming economy, a high level of social organization and advanced metallurgy as well as by large proto-urban mega-sites that may have housed hundreds or thousands of inhabitants during the Middle period (4100-3600 BCE). However, subsequently these settlements were mostly abandoned^4^, and there is archaeological evidence that individuals of the Late CTC interacted with populations that lived in the vast grasslands, or steppes, of Eurasia, such as the Early Bronze Age Yamnaya pastoralists^7^.

**Figure 1.**
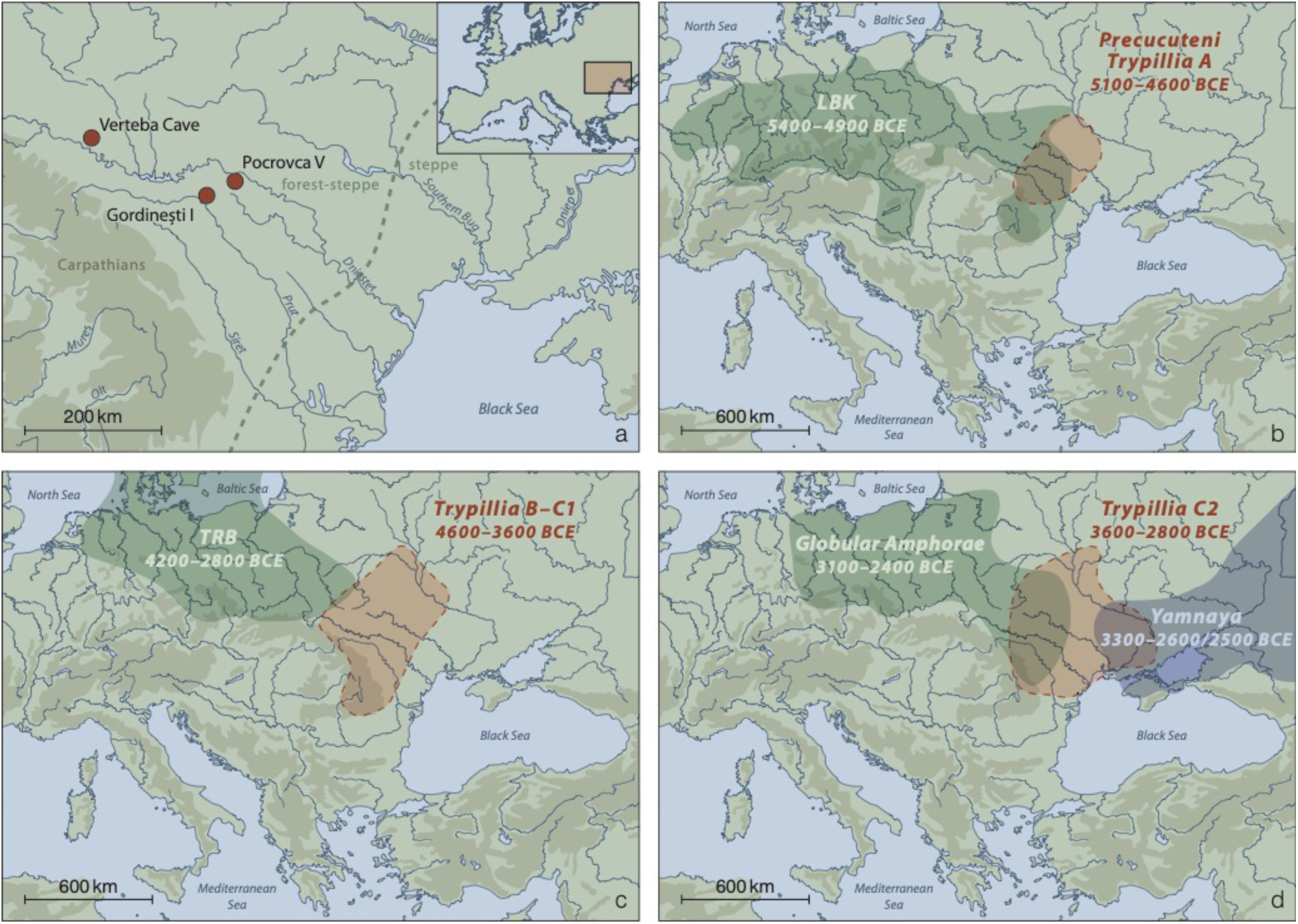
a) Map with the Moldovan sites Pocrovca V and Gordinești I, from where the individuals presented in this study were recovered. Also shown is Verteba Cave in Ukraine, where the CTC individuals presented in Mathieson et al. 2018 were discovered. **b-d)** Temporal and geographic distribution of archaeological cultures mentioned in this study shown for **b)** Early (5100-4600 BCE), **c)** Middle (4600-3600 BCE) and **d)** Late (3600-2800 BCE) period of the CTC.

Despite the important role of CTC in prehistoric eastern Europe, little is known about the genetic composition of the people associated with this complex, the extent of biological contacts with their neighbors or the level of continuity with successive cultural groups. This gap in the genetic landscape can be attributed to a remarkable lack of CTC burials. Human remains have mainly been recovered from Late CTC contexts^6^, and ancient DNA (aDNA) studies have so far only been performed on specimens from a Trypillian site called Verteba Cave in Ukraine (Figure 1). A mitochondrial DNA (mtDNA) study on eight Verteba individuals (3700-2900 cal. BCE) revealed in six cases maternal lineages typical of Anatolian and central European farmers. Two specimens had haplogroup U8b1 that may have been derived from European hunter-gatherers^6, 8^. A subsequent genome-wide analysis in four males from Verteba (3900-3600 cal. BCE) confirmed the large Neolithic (~80%) and smaller hunter-gatherer (~20%) ancestry components^9^.

Here, we present genome-wide data generated from the human remains of four females that were excavated from two Late CTC burials in the present-day Republic of Moldova (Figure 1). The specimens dated to 3500-3100 cal. BCE, i.e. several hundred years later than the previously investigated males from Verteba Cave^9^. The incorporation of these additional data sets obtained from new specimens, sites and time points allows us to draw a more nuanced picture of the population movements and dynamics during this important period in the prehistory of eastern Europe.

## Results

We generated genome-wide shotgun sequencing data from three adults that were recovered from a multiple burial at the site of Pocrovca V (individuals: Pocrovca 1, Pocrovca 2, Pocrovca 3) and from a child interred in the Gordinești I flat necropolis (individual: Gordinești) in northern Moldova (Figure 1). The specimens date to the Late CTC period (3500-3100 cal. BCE). Bioinformatic data analyses showed that all four individuals were females and carried the mitochondrial haplogroups U4, K1, T1 and T2, respectively (Table 1). Kinship analyses revealed no relatedness among them. When we screened the sequence data for known human blood-borne pathogens such as *Yersinia pestis*, *Mycobacterium tuberculosis* and *Mycobacterium leprae*, no signs of an infection were detected.

**Table 1:** Summary information of the Moldova Late Eneolithic CTC samples analyzed in this study. Shown are the archaeological sites, anthropological data, radiocarbon dates, number of SNPs obtained from each sample, genetically determined sex and mitochondrial DNA haplogroup.

| **Sample** | **Anthropological data** | **Calibrated years BCE** | **Called SNPs** | **Genetic sex** | **mtDNA haplogroup** |
| --- | --- | --- | --- | --- | --- |
| Gordinești | child 9 yo | 3482-3197 | 909828 | F | U4a1 |
| Pocrovca 1 | woman 20-25 yo | 3364-3138 | 78116 | F | K1a1 |
| Pocrovca 2 | woman 35-40 yo | 3366-3135 | 1099753 | F | T2c1d1 |
| Pocrovca 3 | woman 60-65 yo | 3341-3114 | 1121523 | F | T1a |

“yo”: years old

A principal component analysis of the four Moldova females together with previously published data sets of ancient Eurasians^9, 10, 11 12, 13^ showed that Gordinești, Pocrovca 1 and Pocrovca 3 grouped with later dating Bell Beakers from Germany and Hungary close to the four CTC males from Verteba, while Pocrovca 2 fell into the LBK cluster next to Neolithic farmers from Anatolia and Starčevo individuals (Figure 2, Supplementary Figure 1).

**Figure 2.**
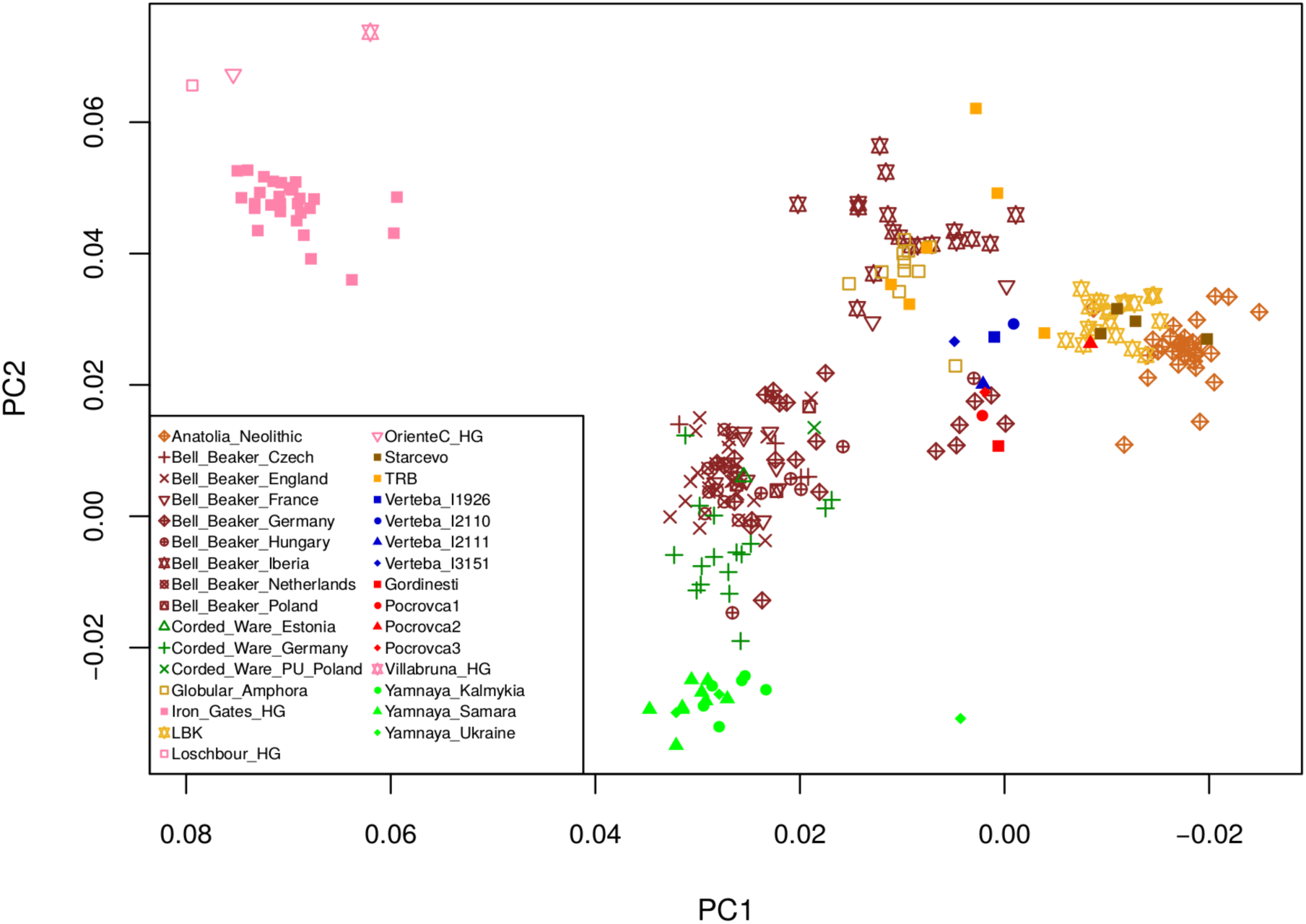
Principal component analysis of the CTC individuals from Moldova (Gordinești, Pocrovca 1, Pocrovca 2, Pocrovca 3) in red and the CTC individuals from Verteba Cave (I1926, I2110, I2111, I3151) in blue together with 23 selected ancient populations/individuals projected onto a basemap of 58 modern-day West Eurasian populations (not shown). HG=hunter-gatherer, LBK=Linearbandkeramik, PU=Proto-Unetice, TRB=Trichterbecher (Funnel Beaker Culture, FBC). PC1 is shown on the x-axis and PC2 on the y-axis.

To assess the genetic diversity within the Moldovan CTC individuals, we ran ADMIXTURE analysis^14^. The dominant element in all Moldovan CTC females was found in Anatolian Neolithic farmers, Starčevo and LBK individuals (Figure 3, Supplementary Figure 2), followed by a large hunter-gatherer component. Interestingly, Gordinești, Pocrovca 1 and 3 had a considerable amount of steppe ancestry, with the Gordinești child exhibiting the highest proportion.

We next applied *f3* outgroup statistics^15^ to investigate which ancient populations or individuals shared most of the genetic drift with the CTC Moldovans. All of them, except Pocrovca 2, appeared genetically close to the Verteba CTC males (Figure 4, Supplementary Figure 3). In contrast, Pocrovca 2 shared most of its genetic ancestry with Starčevo individuals.

**Figure 3.**
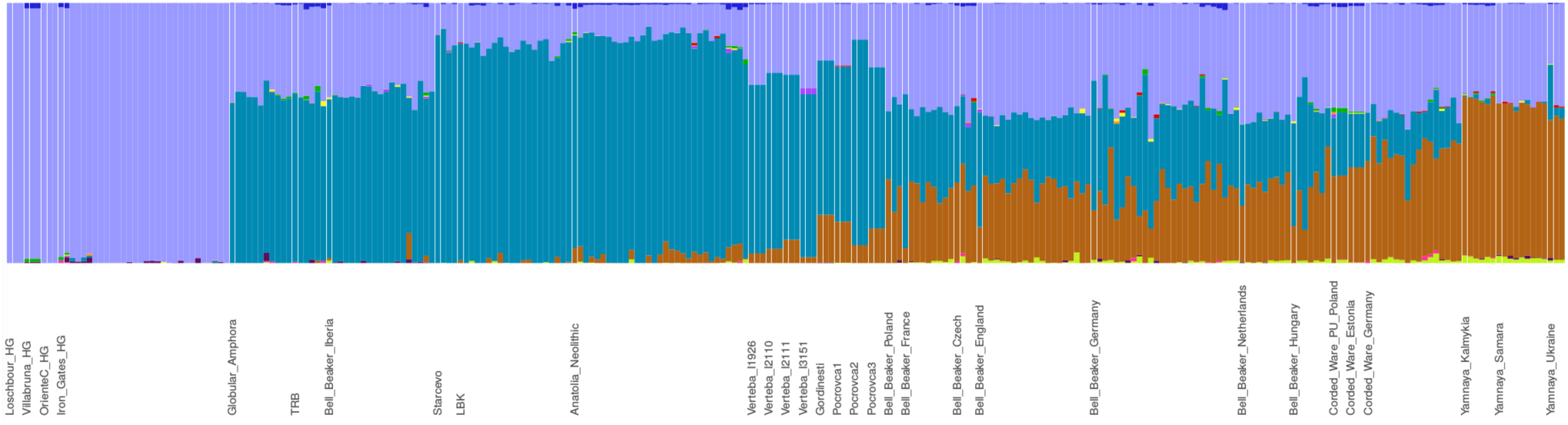
Admixture analysis of the CTC individuals from Moldova and Ukraine together with 23 selected ancient populations/individuals. Admixture plot is shown for K=12 ancestral genetic components. HG=hunter-gatherer, LBK=Linearbandkeramik, PU=Proto-Unetice, TRB=Trichterbecher (Funnel Beaker Culture, FBC). Farmer ancestry is illustrated in turquoise, hunter-gatherer ancestry is shown in purple and steppe-ancestry in brown.

**Figure 4.**
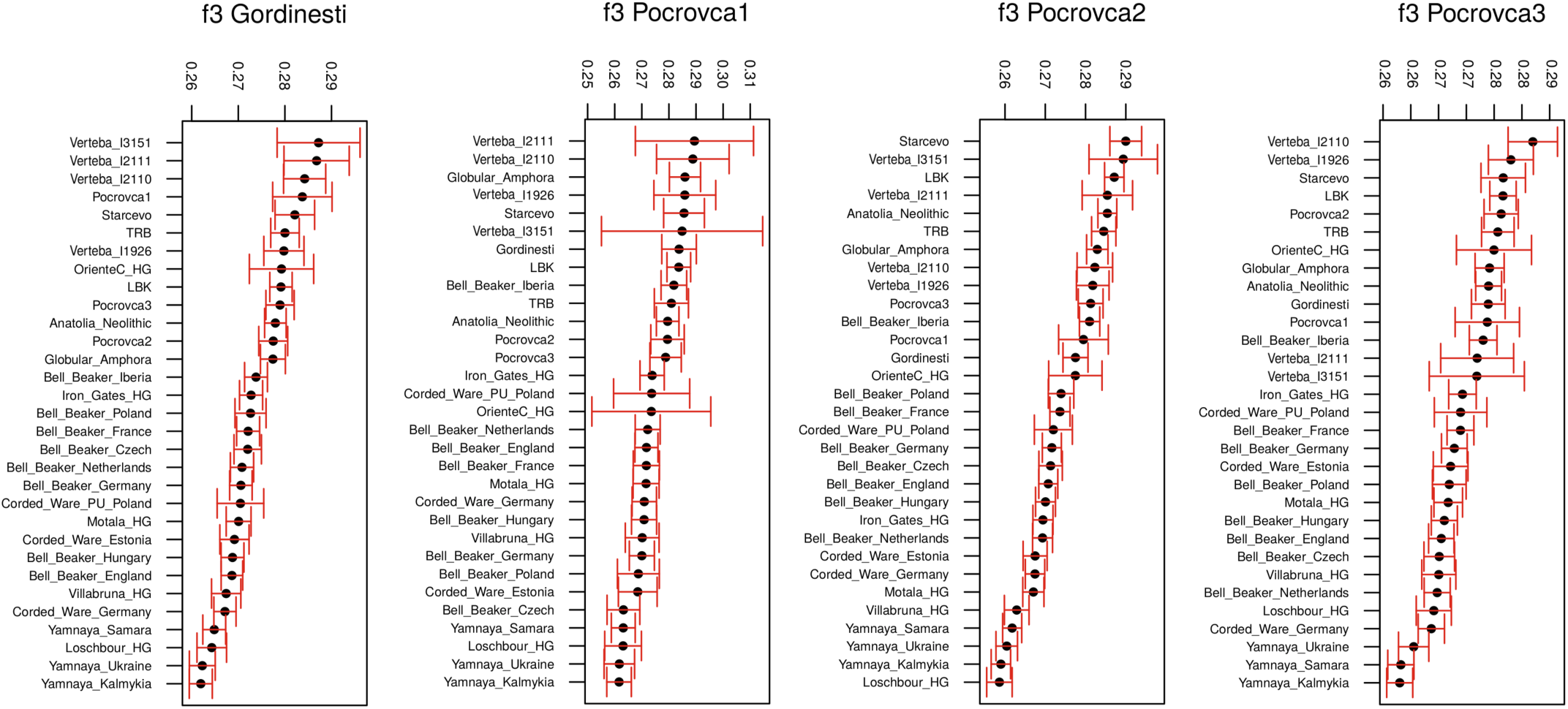
f3 outgroup statistics *f3(Gordinești; Test, Mbuti)* and *f3(Pocrovca; Test, Mbuti)* showing the amount of shared genetic drift between each of the four individuals analysed in this study and selected previously published ancient populations/individuals (as used in PCA and admixture analysis). HG=hunter-gatherer, LBK=Linearbandkeramik, PU=Proto-Unetice, TRB=Trichterbecher (Funnel Beaker Culture, FBC).

We ran D statistics and qpAdm^15^ on the four Moldovan data sets to estimate the direction of genetic influx and the amount of the ancestry components. When the data sets from all four Moldovan CTC individuals were combined, they showed a stronger influx 1) from LBK than Anatolian Neolithic and 2) from Western hunter-gatherers than steppe-related populations. When looking at various proxies for steppe-related ancestry (Yamnaya Samara, Ukraine Mesolithic, Caucasian hunter-gatherer (CHG), Eastern hunter-gatherer (EHG)), we did not observe any significant difference in genetic influx from either Yamnaya Samara, EHG or Ukraine Mesolithic. However, relative to CHG, we detected a substantial shift towards Yamnaya Samara steppe-related ancestry (Supplementary Table 1). Consequently, Yamnaya Samara, Ukraine Mesolithic and EHG appear to be equally suitable proxies for steppe-related ancestry in the Moldovan CTC individuals. This finding was confirmed by our results from the two-way admixture qpAdm models (Supplementary Table 2) for each individual separately and for the combined data.

Next, we applied a three-way admixture qpAdm model to the combined data set of our four Moldovan CTC individuals using LBK, steppe (Yamnaya Samara, EHG, Ukraine Mesolithic) as well as WHG as possible source populations, but did not obtain feasible results. We then modeled each individual separately. Interestingly, Pocrovca 1 yielded a feasible three-way admixture model suggesting the following proportions: LBK (41-60%), steppe-related ancestry (8-18%) and WHG (29-41%) (Supplementary Table 2). This finding is in accordance with the results from our admixture analysis and the D statistics. For the three individuals Gordinești, Pocrovca 2 and Pocrovca 3, the three-way admixture models were not feasible most likely due to an already achieved saturation of the hunter-gatherer ancestry in the steppe populations tested. We did not obtain feasible models when running qpAdm on the X-chromosome in order to test for male-biased admixture from hunter-gatherers or individuals with steppe-related ancestry.

## Discussion

CTC plays an important role in eastern European prehistory, yet owing to the scarcity of human skeletal remains and corresponding aDNA studies, the current knowledge on the origin and biological relatedness of the people associated with this cultural complex is rather limited. Up to now, genome-wide investigations have focused on only four male individuals from a single Trypillian site, Verteba Cave in Ukraine^9^. The vast majority of the human remains found there constituted crania and mandibles that were disposed of in the form of secondary interments. The different bone elements were found commingled and showed high levels of perimortem trauma and postmortem manipulation^16, 17^. In contrast, the skeletal remains from the females presented in this study did not show any signs of interpersonal violence. The two sites Pocrovca and Gordinești, from which the individuals were recovered, are in close proximity to each other and a few hundred kilometers away from Verteba (Figure 1). It should be taken into account that all three locations are situated in the Prut – Dniester interfluve corresponding to the core area of Gordinești local group of Tripolye C2 period or western part of the CTC areal.

Recently, it was hypothesized that due to their high population densities, the CTC mega-settlements served as a focus point for the emergence and large-scale radiation of *Y. pestis* lineages across Eurasia during the Neolithic^18^. Amongst the four Moldovan specimens, we did not detect any signals of a *Y. pestis* infection, although the three individuals from Pocrovca were discovered in a multiple burial (without any traces of violence), which would render death due to an epidemic event plausible.

The genome-wide data sets of the four female individuals presented in this study showed genetic ancestry common in Anatolian farmers and LBK individuals, steppe-related populations as well as Western hunter-gatherers. With maximum 60% the Neolithic-derived proportion constituted the largest ancestry component. Interestingly, from our results the CTC Moldovans appeared to be genetically more likely related to LBK than to Anatolian Neolithic farmers. In the Verteba individuals, the Neolithic component was also seen in the same magnitude but had rather a northwestern Anatolian origin^9^. The high amount of shared genetic ancestry between CTC and LBK or Starčevo, respectively, as observed in our f3-outgroup analysis, is supported by archaeological evidence. The basis for the economic subsistence and cultural attributes of the CTC is found in the European Boian and Starčevo cultures with additional influence from the LBK^19, 20, 21^.

Interestingly, we detected steppe-related ancestry in the Late Eneolithic CTC burials from the Republic of Moldova. The presence of this component suggests moderate genetic influx from individuals affiliated with steppe cultures into the CTC-associated gene-pool as early as 3500 BCE; at the same time, archaeological evidence display an increase of quantity of Tripolye-related finds in the steppe area^22, 23^. Thus, the steppe component had arrived in the eastern part of the continent in farmer communities well before it first appeared in the west, i.e. in the Corded Ware people around 2800 BCE^9^. This finding establishes eastern Europe as an old genetic contact zone between locals and incoming steppe people, which is supported by two other early dating specimens from Ukraine (from Alexandria 4045-3974 cal. BCE and Dereivka 3634-3377 cal. BCE) that also showed mixtures of steppe-and Anatolian Neolithic-related ancestry^9^. However, they represented individuals who still followed a hunter-gatherer subsistence, whereas the CTC females analyzed here belonged to a Neolithic agrarian culture. One likely source population that could have introduced the steppe ancestry component into the CTC gene-pool might have been individuals associated with the eastern Eurasian Mesolithic, e.g. the Ukraine Mesolithic people, Eastern hunter-gatherers or even later-dating Yamnaya steppe pastoralists (Supplementary Table 2). Support for a mixture of Late CTC with the neighboring Early Bronze Age Yamnaya culture exists in the archaeological record (Figure 1; 3300-2600 BCE)^24^. Despite the short time of overlap, artifacts found in both late CTC and Yamnaya settlements provide evidence of barter between them^24^. These observations and the genetic findings presented in this study (i.e. the different steppe proportions in the four Moldovans) are in agreement with ongoing contacts as well as with gradual admixture and a slow change in cultural expression, rather than total replacement. However, this hypothesis challenges a previously published scenario of Yamnaya horsemen massively migrating in war into central Europe^25^.

It is not surprising that Gordinești, Pocrovca 1 and Pocrovca 3 showed genetic affinities with later dating Bronze Age or Bell Beaker individuals. The common link among them is the considerable steppe-related ancestry, which each group likely received independently from different parental populations^26^.

Our analyses (PCA, f3-outgroup analysis) also suggests a genetic relationship between the individuals from Moldova and those associated with the contemporaneous FBC/TRB and GAC, possibly indicating a common origin and/or ongoing interactions. The mtDNA study on the Verteba individuals already showed a high degree of similarity in the maternal lineage composition between CTC and FBC populations^6^. This connection can be explained by the geographical proximity of the FBC and CTC distribution areas (Figure 1). An overlap of CTC and FBC settlements site has been documented and there is additional confirmation in the archaeological record for regular inter-group contacts and trade westwards and northwards from the CTC into the GAC and FBC areas^27^.

Overall, the different genetic makeup of the CTC individuals presented here and in Mathieson et al. 2018 indicates a relatively high diversity, which is surprising given that they all dated to the same Late CTC period and were buried only a few hundred kilometers apart (Figure 1). This finding suggests population dynamics also within a culture and questions the notion of the apparently stable and uniform composition of individuals associated with a specific archaeological group.

## Material and Methods

### Samples

Pocrovca V: the collective burial belonging to the Trypillia C2 culture (Gordinești local group) contained three human skeletons as well as pottery and animal bones including those of three hares. The bodies of the deceased were thrown into the pit sequentially, one on top of the other. They lay in a crouched position on their right sides, which is the main inhumation type for this cultural group. Osteologically, the remains represented adult females between 20 and 65 years old (Table 1, detailed description SI).

Gordinești: the grave in the Gordinești I flat necropolis belonging to the Trypillia C2 culture (Gordinești local group) contained the incomplete skeleton of a child. The age at death estimation of 9 years ± 24 months was based on the dentition (detailed description SI).

All of the specimens were radiocarbon-dated to 3500-3100 cal. BCE (Table 1).

### Ancient DNA extraction and library preparation

Petrous bones and teeth were cleaned in pure bleach solution for 5 minutes and rinsed with water. After drying overnight at 37° C the inner ear area (cochlea and vestibule) was cut out from the petrous bone, bleached, rinsed with water and dried. The dried inner ear piece was ground in a ball mill homogenizer for 45 s at maximum speed. Fifty mg of bone powder were used for DNA extraction following a silica-based protocol^28^. For each sample, a double-stranded DNA sequencing library was prepared from 20 µL of extract, following partial uracil-DNA-glycosylase treatment^29^. Sample-specific index combinations were added to the sequencing libraries in order to allow differentiation between the individual samples after pooling and multiplex sequencing^30^. Sampling, aDNA extraction and the preparation of sequencing libraries were performed in clean-room facilities of the Ancient DNA Laboratory in Kiel. Negative controls were taken along for the DNA extraction and library generation steps.

### Sequencing

The libraries were paired-end sequenced using 2×75 cycles on an Illumina HiSeq 4000. Demultiplexing was performed by sorting all the sequences according to their index combinations. Illumina sequencing adapters were removed and paired-end reads were merged. Merged reads were filtered for a minimum length of 30 bp.

### Screening for pathogens

All samples were screened for their metagenomic content with the metagenome analyzer MEGAN^31^ and the alignment tool MALT^32^. MALT version V0.3.8 was used to align all pre-processed samples against a collection of available complete bacterial and viral genomes. Bacterial genomes were downloaded from the NCBI FTP server (ftp.ncbi.nlm.nih.gov/genomes/refseq/bacteria, access 12.03.2018) using a custom script. Viral genomes were downloaded from the NCBI FTP server (ftp://ftp.ncbi.nlm.nih.gov/refseq/release/viral/, access 03.01.2018). MALT was executed in BLASTN mode with the following parameters for bacteria:

*malt-run--mode BlastN-e 0.001-id 95--alignmentType SemiGlobal--index $REF –inFile $IN –output $OUT*

and for viruses:

*malt-run--mode BlastN-e 0.001-id 85--alignmentType SemiGlobal--index $REF –inFile $IN –output $OUT*

where $REF is the MALT index, $IN is a clipped-and-merged FASTQ file and $OUT is the output folder for MALT. Resulting RMA files were examined for their taxonomic content using MEGAN version V6.11.4.

### Mapping and aDNA damage patterns

Pre-processed sequences were mapped to the human genome build hg19 (International Human Genome Sequencing Consortium, 2001) using BWA 0.7.12^33^ with the reduced mapping stringency parameter “-n 0.01” to account for mismatches in aDNA. Duplicates were removed. In order to assess the authenticity of the aDNA fragments^34^, terminal C to T mis-incorporations were evaluated using mapDamage 2.0^35^. After the validation of damage, the first two positions at the 5’ end of the fastq-reads were trimmed off.

### Sex determination

Sex was determined based on the ratio of sequences aligning to the X and Y chromosomes compared to the autosomes^36^. Females are expected to have a ratio of 1 on the X chromosome and 0 on the Y chromosome, whereas males are expected to have both X and Y ratios of 0.5.

### Genotyping

Alleles were drawn at random from each of the 1,233,013 SNP positions^11, 13^ in a pseudo-haploid manner using a custom script as described in Lamnidis et al. 2018^37^. 10,000 SNPs served as a minimum threshold required for a sample to be included in further analyses.

### Contamination estimation and authentication

Estimation of exogenous contamination in the aDNA extracts was performed on the mitochondrial level using the software Schmutzi^38^.

### Kinship analysis

Kin relatedness was assessed among the four Moldova CTC individuals using READ^39^ and lcMLkin^40^. READ identifies relatives based on the proportion of non-matching alleles. lcMLkin infers individual kinship from calculated genotype likelihoods.

### Principal component analysis (PCA)

Genotype data sets of the Moldovan samples were merged with previously published genotypes of 5,514 modern and ancient individuals on a data set of 597,573 SNPs^11, 12, 13, 15^. Using the software Smartpca (version 16000)^41^, the ancient individuals were projected on a basemap of genetic variation calculated from the following 58 West-Eurasian populations^11, 12, 13^: Abkhasian, Adygei, Albanian, Armenian, Balkar, Basque, Bedouin, Belarusian, Bergamo, Bulgarian, Canary Islander, Chechen, Croatian, Cypriot, Czech, Druze, English, Estonian, Finnish, French, Georgian, Greek, Hungarian, Icelandic, Iranian, South Italian, Jewish (Ashkenazi, Georgian, Iranian, Iraqi, Libyan, Moroccan, Tunisian, Turkish, Yemenite), Jordanian, Kumyk, Lebanese, Lezgin, Lithuanian, Maltese, Mordovian, North Ossetian, Norwegian, Orcadian, Palestinian, Russian, Sardinian, Saudi, Scottish, Sicilian, Spanish, North Spanish, Syrian, Turkish, Tuscan, Ukrainian.

### ADMIXTURE analysis

Prior to ADMIXTURE analysis we used Plink (v1.90b3.29) to prune out SNPs in linkage disequilibrium using the parameters *--indep-pairwise 200 25 0.4.* ADMIXTURE (version 1.3.0)^14^ was run on a data set of 597,573 SNPs^11, 12, 13, 15^ comprising 5,514 previously published ancient and modern human samples and our four Moldovan samples. We used a number of ancestral components (K) ranging from 4 to 12 in 100 bootstraps for each component, respectively.

### D statistics

D statistics were run as part of the Admixtools package^15^ in the form of *(Mbuti; Moldova; PopC; PopD)*, iteratively shuffling different proxies for farmer-related ancestry, hunter-gatherer-related ancestry and steppe-related ancestry, such as LBK and Anatolian Neolithic, Western, Eastern and Caucasian hunter-gatherers and Yamnaya Samara for PopC and PopD, respectively.

### f3-outgroup statistics

f3-outgroup statistics were run as a part of the Admixtools package^15^ in the form of *f3 (Moldova; Test; Mbuti)* using for *Test* the same populations as in the ADMIXTURE and PCA analyses.

### qpAdm analysis

qpAdm analysis^15^ was run to model the Moldovan individuals as admixture of farmers, hunter-gatherers or individuals with steppe-related ancestry, such as Yamnaya. The following populations were used as outgroups: Mbuti, Ust Ishim, Kostenki14, MA1, Han, Papuan, Onge, Chukchi and Karitiana.

### Determination of mitochondrial haplogroups

Sequencing reads were mapped to the human mitochondrial genome sequence rCRS^42^. Consensus sequences were generated in Geneious (v. 9.1.3) using a default threshold of 85% identity among the covered positions and a minimum coverage of 3. HAPLOFIND^43^ was applied to determine the mitochondrial haplogroups from the consensus sequences.

### Investigating male-bias in steppe-related ancestry admixture

qpAdm was used to estimate admixture proportions on the autosomes compared to the X-chromosome in order to compute Z-scores for the difference between autosomes and the X-chromosome as described in Mathieson et al. 2018, where a positive Z-score indicates a male-biased admixture through steppe populations, such as Yamnaya.

## Acknowledgment

This study was funded by the Deutsche Forschungsgemeinschaft (DFG, German Research Foundation - Projektnummer 2901391021 – SFB 1266). We are grateful to Wolfgang Haak for his advice. We thank Ralf Opitz for graphics support. We acknowledge financial support by Land Schleswig-Holstein within the funding programme Open Access Publikationsfonds.

## Author Contribution

B.K.-K., A.N., J.M. and S.T. conceived and designed the research. B.K.-K. generated and A.I., J.S. analyzed the ancient DNA data. A.I., S.T., A.S., J.S., O.S., G.S., R.H., J.M. A.N. and B.K.-K. interpreted the findings. A.N., A.I. and B.K.-K. wrote the manuscript with input from S.T., J.M., A.S..

---

**Supplementary Figure 1.** Principal component analysis of the CTC individuals from Moldova (Gordinești, Pocrovca 1, Pocrovca 2, Pocrovca 3) in red and the CTC individuals from Verteba Cave (I1926, I2110, I2111, I3151) in blue together with 73 previously published ancient populations/individuals projected onto a basemap of 58 modern-day West Eurasian populations (not shown). EBA=Early Bronze Age, EN=Eneolithic, HG=hunter-gatherer, IA=Iron Age, LBA=Late Bronze Age, LBK=Linearbandkeramik, LN=Late Neolithic, M(L)BA=Middle (Late) Bronze Age, MN=Middle Neolithic, PU=Proto-Unetice, TRB=Trichterbecher (Funnel Beaker Culture, FBC). PC1 is shown on the x-axis and PC2 on the y-axis.

**Supplementary Figure 2.** Unsupervised admixture analysis for K=4, 6, 8 and 12 ancestral genetic components for 117 selected ancient populations/individuals. Modern-day populations were included in the analysis but are not shown. BA=Bronze Age, EBA=Early Bronze Age, EN=Eneolithic, HG=hunter-gatherer, IA=Iron Age, LBA=Late Bronze Age, LBK=Linearbandkeramik, LN=Late Neolithic, M(L)BA=Middle (Late) Bronze Age, MN=Middle Neolithic, PPNA=Pre-Pottery Neolithic A, PPNB=Pre-Pottery Neolithic B, PU=Proto-Unetice, TRB=Trichterbecher (Funnel Beaker Culture, FBC).

**Supplementary Figure 3.** f3 outgroup statistics *f3(Gordinești; Test, Mbuti)* and *f3(Pocrovca; Test, Mbuti)* showing the amount of shared genetic drift between our CTC individuals from Pocrovca and Gordinești, respectively, and previously published ancient populations/individuals. EBA=Early Bronze Age, EN=Eneolithic, HG=hunter-gatherer, IA=Iron Age, LBA=Late Bronze Age, LBK=Linearbandkeramik, LN=Late Neolithic, M(L)BA=Middle (Late) Bronze Age, MN=Middle Neolithic, PU=Proto-Unetice, TRB=Trichterbecher (Funnel Beaker Culture, FBC).

**Supplementary Table 1.** D statistics for all four Moldovan data sets Gordinești, Pocrovca 1, Pocrovca 2, Pocrovca 3 individually and combined.

**Supplementary Table 2.** qpAdm three-way and two-way admixture modelling for all four Moldovan data sets Gordinești, Pocrovca 1, Pocrovca 2, Pocrovca 3 individually and combined.

